# Inhibitors of spore germination act against diverse human fungal pathogens

**DOI:** 10.64898/2026.09.15.751812

**Authors:** Jacqueline A. Spieles, Seth D. Greengo, Ashley M. Holt, Madison C. Barnes, Nancy P. Keller, Jeniel E. Nett, Christina M. Hull

**Affiliations:** Department of Biomolecular Chemistry, University of Wisconsin, Madison Madison, WI 53706; Department of Medical Microbiology & Immunology, University of Wisconsin, Madison Madison, WI 53706; Department of Medicine, School of Medicine and Public Health, Department of Plant, University of Wisconsin, Madison Madison, WI 53706; Pathology, College of Agricultural and Life Sciences, University of Wisconsin, Madison Madison, WI 53706

**Author notes:** To whom correspondence should be addressed: Dr. Christina M. Hull, 1135 Biochemistry Building, 420 Henry Mall, University of Wisconsin, Madison, Madison, WI 53706. Department of Medicine, Medical College of Wisconsin, 8701 Watertown Plank Road, Milwaukee, WI 53226.

**Keywords:** *Cryptococcus neoformans*, *C. deneoformans*, *Aspergillus fumigatus*, spores, germination, antifungal agents, conidia

## Abstract

Current antifungal drugs often cause severe side effects due to the conservation of molecular targets between fungal and mammalian cells. One way to avoid host toxicity in future antifungals is to leverage molecules with targets involved in processes unique to fungi, such as spore germination. Germination is conserved in several of the most critical fungal pathogens but absent in mammalian cells. In this work we evaluated 191 small molecule inhibitors of *Cryptococcus* germination for their abilities to inhibit germination of another fungal pathogen, *Aspergillus fumigatus*, as well as yeast growth of *Candida albicans* and a pan-resistant isolate of *Candida* (*Candidozyma*) *auris*. The cytotoxicity of these inhibitors towards human hepatocytes and erythrocytes was also assessed. Overall, we identified 101 molecules that were either 1) germination-specific inhibitors in one or both germinating species tested, or 2) inhibitory to both germination and vegetative growth in one or more species tested. The majority of these molecules exhibited minimal toxicity to the human cell types tested. Molecules were scored and ranked for therapeutic potential based on assay data. For two key molecules of interest, predicted binding targets (histone deacetylase and acetyl-CoA carboxylase) were identified computationally. Overall, the inhibitor molecules characterized here showed activity against phylogenetically diverse human fungal pathogens with minimal toxic effects on human cell lines.

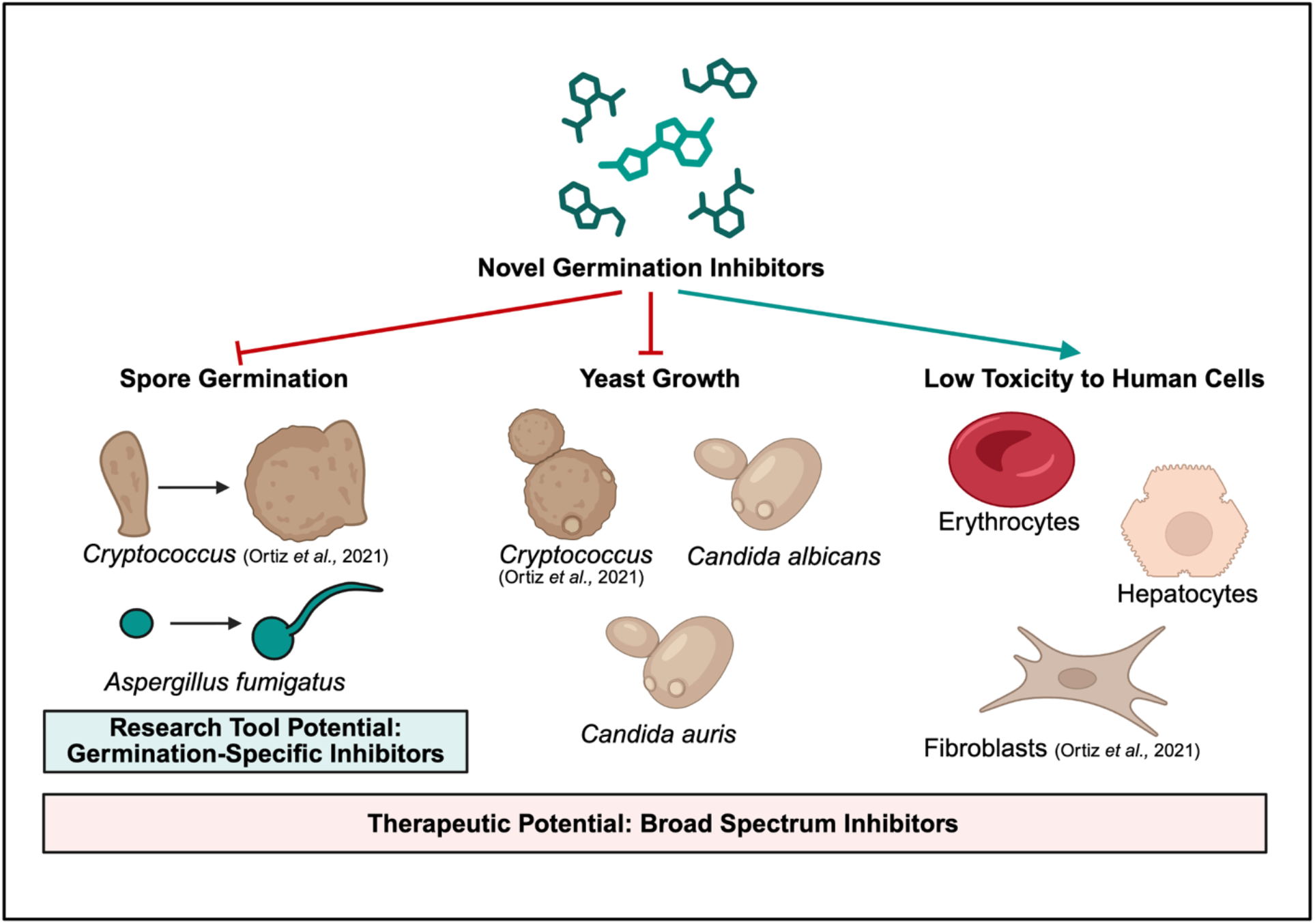

## Introduction

Invasive fungal diseases have emerged in recent decades as major causes of morbidity and mortality worldwide, especially in immunocompromised populations. Even with the administration of antifungal therapies, mortality rates for invasive mycoses can reach upwards of 50% (Denning, 2024). Current antifungal therapies can also induce severe side effects in patients and are increasingly susceptible to antimicrobial resistance (Van Rhijn & White, 2025). Antifungal drug development is challenging due to the shared eukaryotic ancestry between fungi and humans, which limits molecular opportunities to selectively target fungal cells without causing toxicity to the host. One way to address the need for improved antifungal therapeutics is to identify new fungal-specific molecular processes that could be chemically targeted, ideally with a compound that has broad-spectrum activity against multiple fungal pathogens, no/low toxic effects on mammalian cells, and a low risk of resistance development.

Of the 19 fungal pathogens on the World Health Organization Fungal Priority Pathogens List, 13 of them are known to produce spores (WHO FPPL, Gauthier *et al*., 2015, Guarner *et al*., 2011). Spores are a non-replicating, stable, and stress-resistant cell type generally produced when local conditions are not conducive to growth. Fungal pathogens produce spores in environmental reservoirs that can be aerosolized and dispersed to new environments. After dispersal, spore germination into a vegetatively growing cell type is ultimately required for reproduction and survival of the species. For human fungal pathogens, the ability to germinate in the mammalian host is a key requirement for disseminated disease to occur (Ernst *et* al., 2000, Gauthier *et al*., 2015, Guarner *et al*., 2011). Because germination appears to be a fungal-specific process not shared by mammalian host cells and is required for invasive disease, it may harbor novel targets for antifungal drug development. Inhibitors of spore germination could be used prophylactically in populations at greatest risk to prevent invasive fungal disease (Köhler *et al*., 2015, Fisher *et al*., 2012).

Efforts to identify fungal spore germination inhibitors include a high throughput screen of small, drug-like molecules that identified 191 novel inhibitors of *Cryptococcus deneoformans* spores. *C. deneoformans* and its close relative, *Cryptococcus neoformans* are the most common causes of fungal meningoencephalitis and cause upwards of 147,000 deaths per year worldwide (Denning, 2024). *C. deneoformans* has been developed as a model for spore studies because its spores can be readily purified in large quantities for comprehensive studies (Ortiz *et al*., 2021).

The majority of the *C. deneoformans* germination inhibitors identified also inhibit *C. deneoformans* yeast growth and exhibit low cytotoxicity against mammalian fibroblasts (Ortiz *et al*., 2021). To evaluate the efficacy of these compounds in the inhibition of other human fungal pathogens, we carried out quantitative, microscopy-based germination assays using conidia from *Aspergillus fumigatus* and vegetative growth assays using *Candida* (*Candidozyma*) *auris* and *Candida albicans* yeast. *A. fumigatus* is a filamentous ascomycete and the primary cause of pulmonary aspergillosis, which causes an estimated 1.8 million annual deaths globally (Denning, 2024). *C. albicans* and *C. auris* are two of the most prominent causative agents of invasive candidiasis, with *C. auris* being of particular concern due to its high rate of antifungal drug resistance (Pappas *et al*., 2018). These fungal species belong to two phyla that diverged ~450 million years ago: *Ascomycota* (*Candida* and *Aspergillus* species) and *Basidiomycota* (*Cryptococcus* species) (Berbee *et al*., 2010) **(Fig. 1)**. We discovered that despite this large phylogenetic distance between species, a majority of the inhibitors of *C. deneoformans* basidiospore germination were effective against *A. fumigatus* conidia germination, implying that fungal spore germination mechanisms may be more conserved across phylogenies than previously thought. Importantly, most of the compounds were not toxic to human cells. Overall, we have identified broad-spectrum antifungal agents that can be used as research tools to study spore germination and leveraged in the development of antifungal therapeutics.

**Figure 1.**
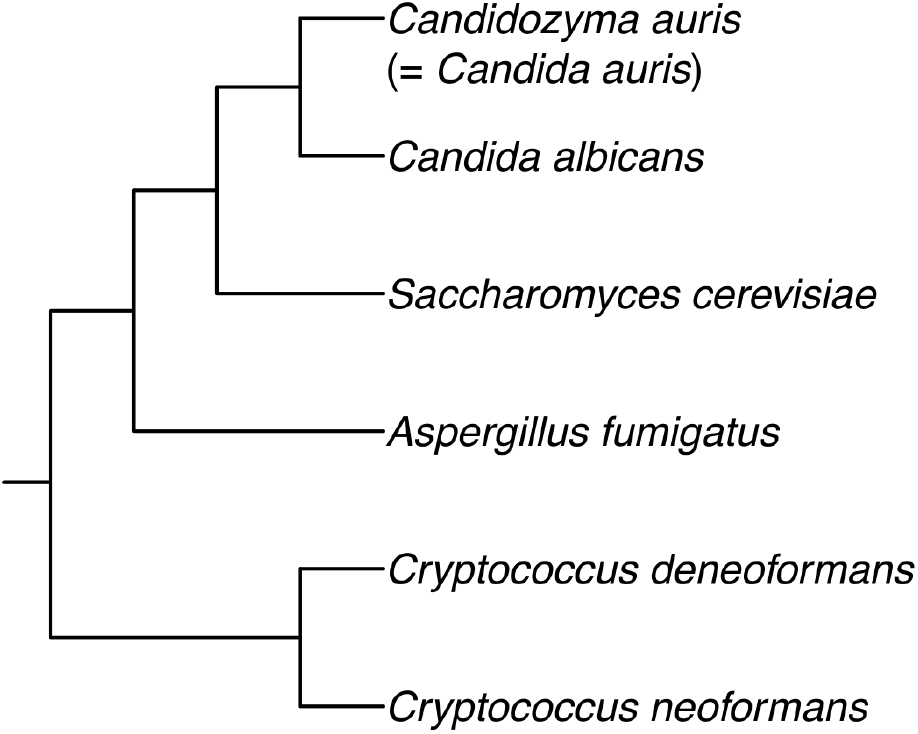
Cladogram illustrating the phylogenetic relationships among the *Candida, Aspergillus*, and *Cryptococcus* species used for inhibitor screening in this work, as well as a nonpathogenic *S. cerevisiae* reference species. Cladogram is based on Li *et al*., 2021 and Shen *et al*., 2016 and was generated using the R packages ggtree v3.14.0 (Yu *et al*., 2017) and ggplot2 v4.0.0 (Wickham, 2009) in R v4.4.2 (R Core Team 2024).

## Results

### 101 *Cryptococcus* spore germination inhibitors also inhibited *Aspergillus* conidia germination

To determine whether 191 previously identified *C. deneoformans* germination inhibitors were also inhibitory to other phylogenetically diverse fungal pathogens, an *Aspergillus*-adapted quantitative germination assay (QGA) was first optimized for screening (Ortiz *et al*., 2021; Ortiz *et al*., 2025) **(Fig. 1)**. QGAs quantify germination progression by measuring changes in cell area (µm^2^) and aspect ratio (width/length) of individual cells in a population over time **(Fig. 2)**. In *Aspergillus*, conidia begin as relatively small, round cells that swell during early germination before extending germ tubes and transitioning into larger, more oblong germlings over approximately eight hours in these assay conditions. Conidia from *A. fumigatus* strain CEA10 were germinated in RPMI-MOPS medium for a total of eight hours at 37 °C and photographed every two hours. Approximately 16,000 conidia per well were evaluated across two wells at three positions per well for a total assessment of approximately 6,000 conidia per inhibitor.

**Figure 2.**
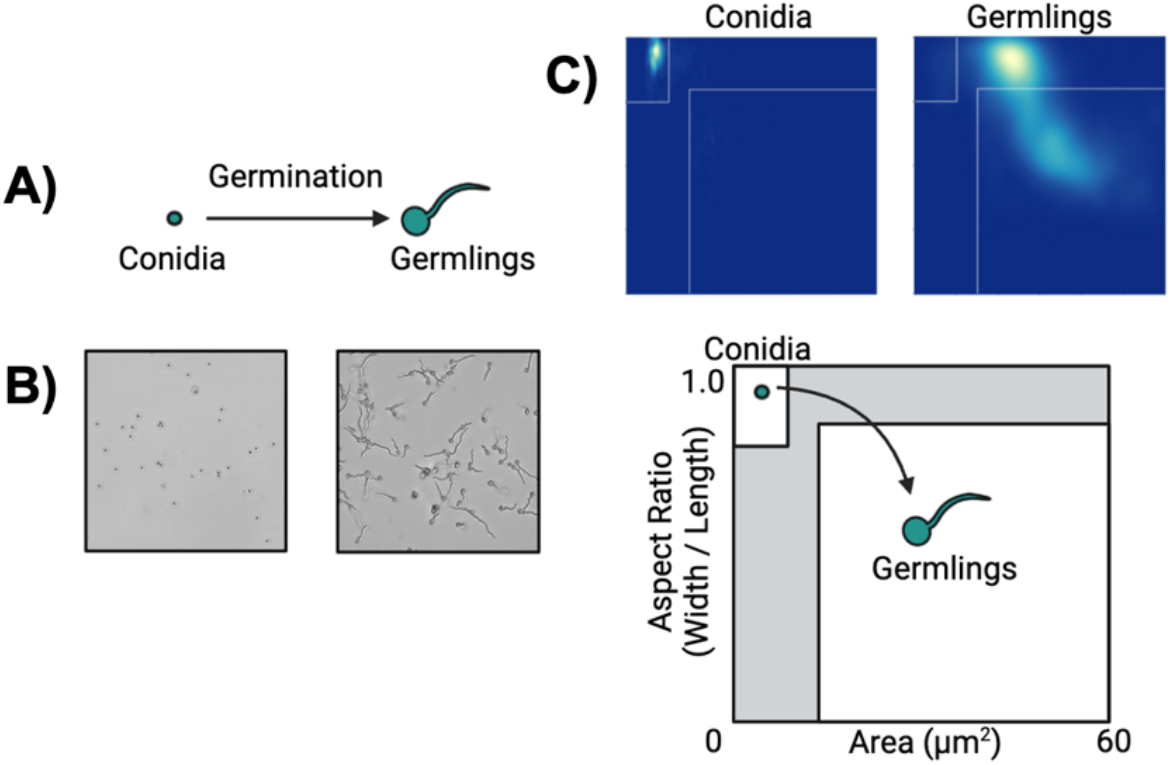
A quantitative germination assay (QGA) adapted for *Aspergillus fumigatus* was used to identify small molecule germination inhibitors (Ortiz *et al*., 2024). **A)** Illustration of *Aspergillus* germination. **B)** Populations of germinating conidia in 384 well plates were microscopically imaged, and **C)** an ImageJ-based analysis pipeline was used to measure aspect ratio (width/length) and area (μm^2^) as conidia germinated into germlings in RPMI-MOPS at 37 °C over 8 hours. Small, round conidia (top left quadrant) germinate into larger, more oblong germlings (bottom right quadrant). Pixel intensity in two-dimensional histograms is representative of relative number of cells per pixel. ~6,000 cells imaged per histogram panel.

We identified 101 compounds that inhibited *A. fumigatus* germination from a total of 191 *C. deneoformans* germination inhibitors. Screening was first conducted in duplicate at 10 µM. Compounds that did not inhibit at 10 µM were retested in duplicate at 100 µM. Of the 120 compounds that did not initially inhibit *A. fumigatus* germination at 10 µM, 30 of them inhibited germination at 100 µM, resulting in a total of 101 small molecules that inhibit germination of both *C. deneoformans* and *A. fumigatus* spores **(Fig. 3, Dataset S1)**.

**Figure 3.**
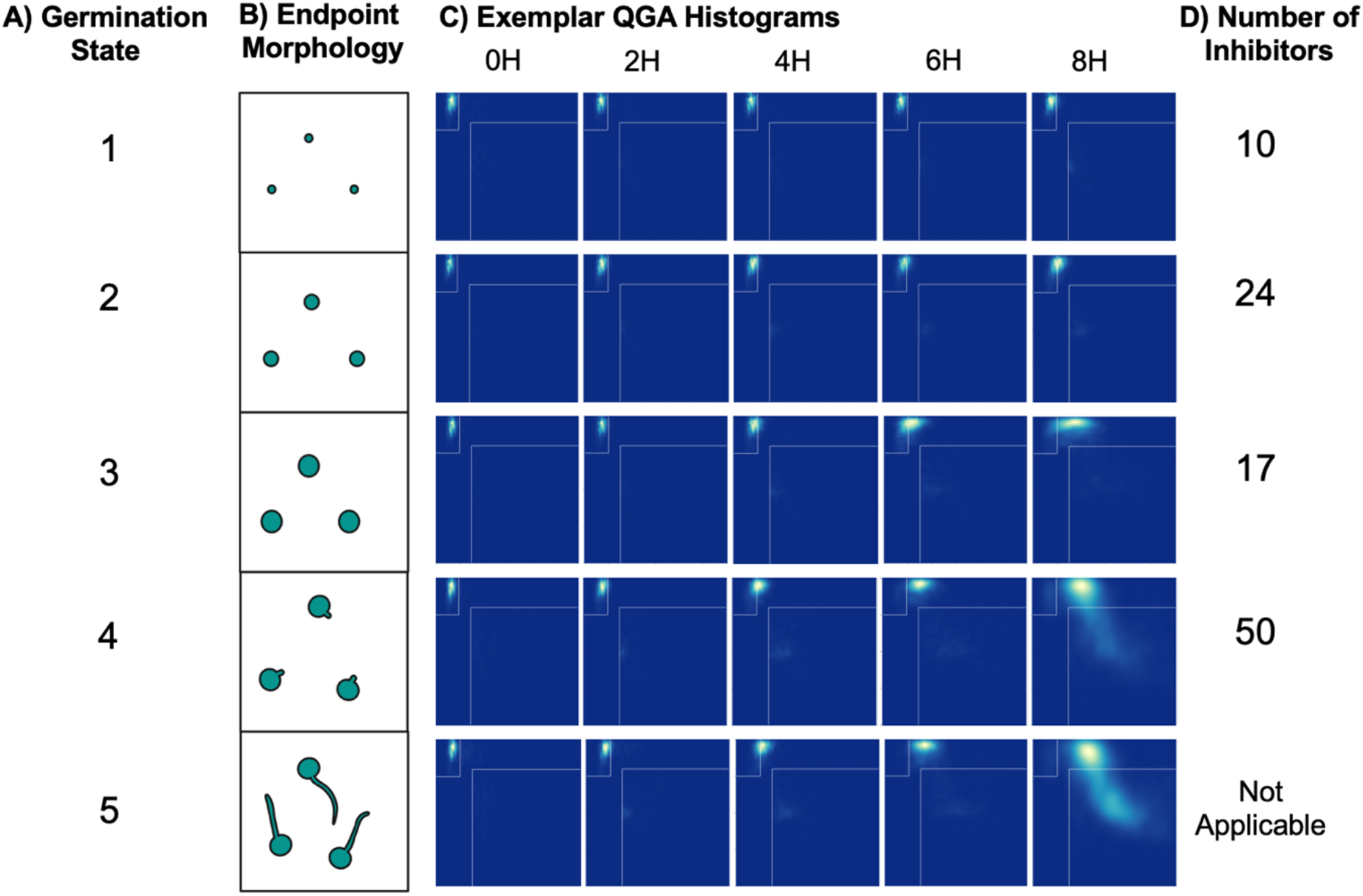
Of 191 previously identified *C. deneoformans* germination inhibitors, 101 also inhibited germination of *A. fumigatus*. **A)** Inhibition was parsed into Germination States 1-5, based on endpoint morphology, where 1 is complete inhibition of morphological changes, and 5 is no inhibition (complete germination into germlings). **B)** Illustrated cell morphologies at the eight-hour endpoint for each Germination State. **C)** QGA 2D histograms for exemplar inhibitors belonging to each Germination State. **D)** Number of inhibitors belonging to each Germination State. Experiments were conducted with compounds in duplicate wells.

At the eight-hour experimental endpoint, a range of population-level phenotypes was identified and classified into five groups defined as “Germination States” **(Fig. 3A)**. Compounds were identified as inhibitors if they were assigned to Germination States 1-4. Germination State 1 reflected strong inhibition characterized by a complete lack of cell morphology changes associated with germination (indistinguishable from ungerminated conidia). Germination State 2 was marked by partial conidial swelling, Germination State 3 by a stall after complete swelling, Germination State 4 by partial germ tube formation, and Germination State 5 by full, uninhibited germination comparable to no-inhibitor (DMSO-only) controls **(Fig. 3B, 3C)**. These endpoint phenotypes may reflect differences in inhibitor potencies, the abilities of compounds to access intracellular targets through the *A. fumigatus* cell wall, or differences in molecular targets.

Most inhibitors displayed relatively weak or late activity during germination, although several compounds exhibited stronger or earlier inhibitory effects. Ten of the 101 inhibitors were classified as Germination State 1 and fully prevented detectable morphological changes associated with germination by eight hours. Twenty-four inhibitors produced Germination State 2 phenotypes, 17 resulted in Germination State 3 phenotypes, and the remaining 50 inhibitors resulted in Germination State 4 phenotypes, reflecting the weakest or latest inhibition **(Fig. 3D)**.

Additionally, inhibitors of both *A. fumigatus* and *C. deneoformans* germination belonged to shared chemical substructure groups. Of 191 compounds identified previously as *C. deneoformans* germination inhibitors, 76 could be sorted into one of eight chemical substructure groups. In general, compounds in the same substructure group elicited the same inhibition phenotype, suggesting that similarly structured compounds share a common cellular target (Ortiz *et al*., 2021). Fifty-four of these 76 compounds also inhibited *A. fumigatus* germination, with representation in every substructure group **(Table 1, Table S1)**. These findings suggest that inhibitory compounds with shared substructures act on the same target in both fungi and point to conserved molecular mechanisms of germination for these phylogenetically diverged organisms.

**Table 1.**
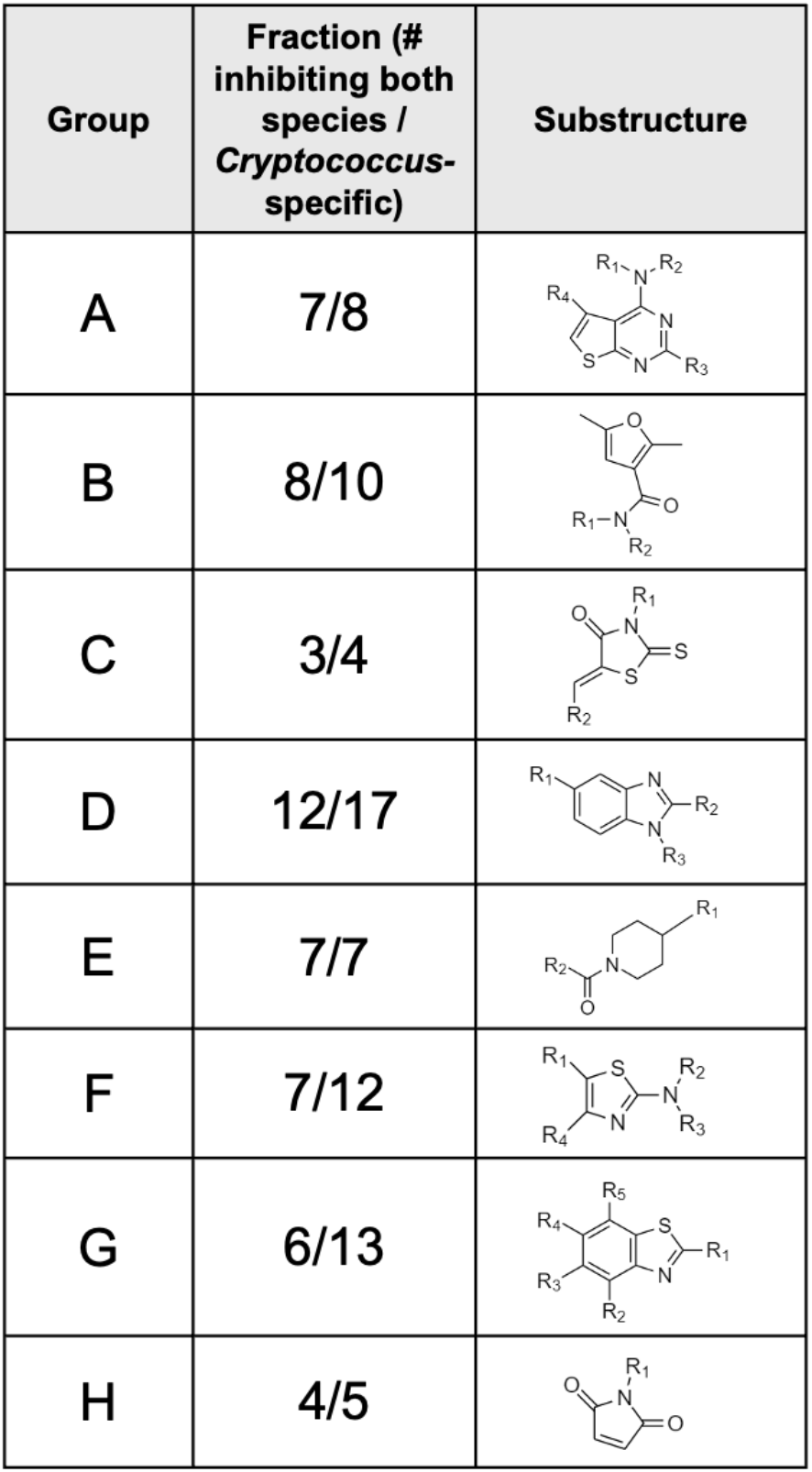
Fifty-four of 101 identified *A. fumigatus* germination inhibitors belong to eight substructure groups defined in a previous screen for *C. deneoformans* germination inhibitors (Ortiz *et al*., 2021). Diagram of each substructure group and fraction indicating number of compounds capable of also inhibiting *A. fumigatus* germination at or below 100 μM.

Both cross-species and species-specific germination inhibitors were identified
| Group | Fraction (# inhibiting both species / *Cryptococcus*-specific) | Substructure |
| --- | --- | --- |
| A | 7/8 |  |
| B | 8/10 |  |
| C | 3/4 |  |
| D | 12/17 |  |
| E | 7/7 |  |
| F | 7/12 |  |
| G | 6/13 |  |
| H | 4/5 |  |

Together, these results indicate that most shared inhibitors act after conidial swelling but before full germ tube formation in *A. fumigatus* and that a substantial fraction of *C. deneoformans* germination inhibitors also impair *A. fumigatus* germination, showing a high level of cross-species activity.

### Both cross-species and species-specific germination inhibitors were identified

Fungal spore germination is a distinct process from vegetative growth. This distinction is important to consider both in the context of infection and basic fungal biology. Of the original 191 *C. deneoformans* germination inhibitors identified, 152 also inhibit at least 25% of *C. deneoformans* yeast growth (Ortiz *et al*., 2021). The remaining 39 molecules inhibit germination specifically. We hypothesized that at least some of these germination-specific inhibitors have targets conserved across phylogenies and would also inhibit *A. fumigatus* germination. We identified 13 molecules that significantly inhibit both *C. deneoformans* and *A. fumigatus* germination but exhibit limited inhibitory activity on *C. deneoformans* yeast growth (over 90% *C. deneoformans* germination inhibited, Germination State 4 or lower inhibition of *A. fumigatus*, and under 25% *C. deneoformans* yeast growth inhibited). Two of these, 71797126 and 27460748 (PubChem CIDs) were among the most extreme examples with the highest relative germination inhibition in both species and the lowest relative yeast growth inhibition. **(Fig. 4A)**. Minimal information is available on either of these compounds as they tested negative for activity in all reported assays in the PubChem bioactivity database, which broadly assesses activity against cellular machinery in bacterial pathogens. It is possible, therefore, that these molecules could aid in characterization of processes unique to and conserved among eukaryotic microbes.

**Figure 4.**
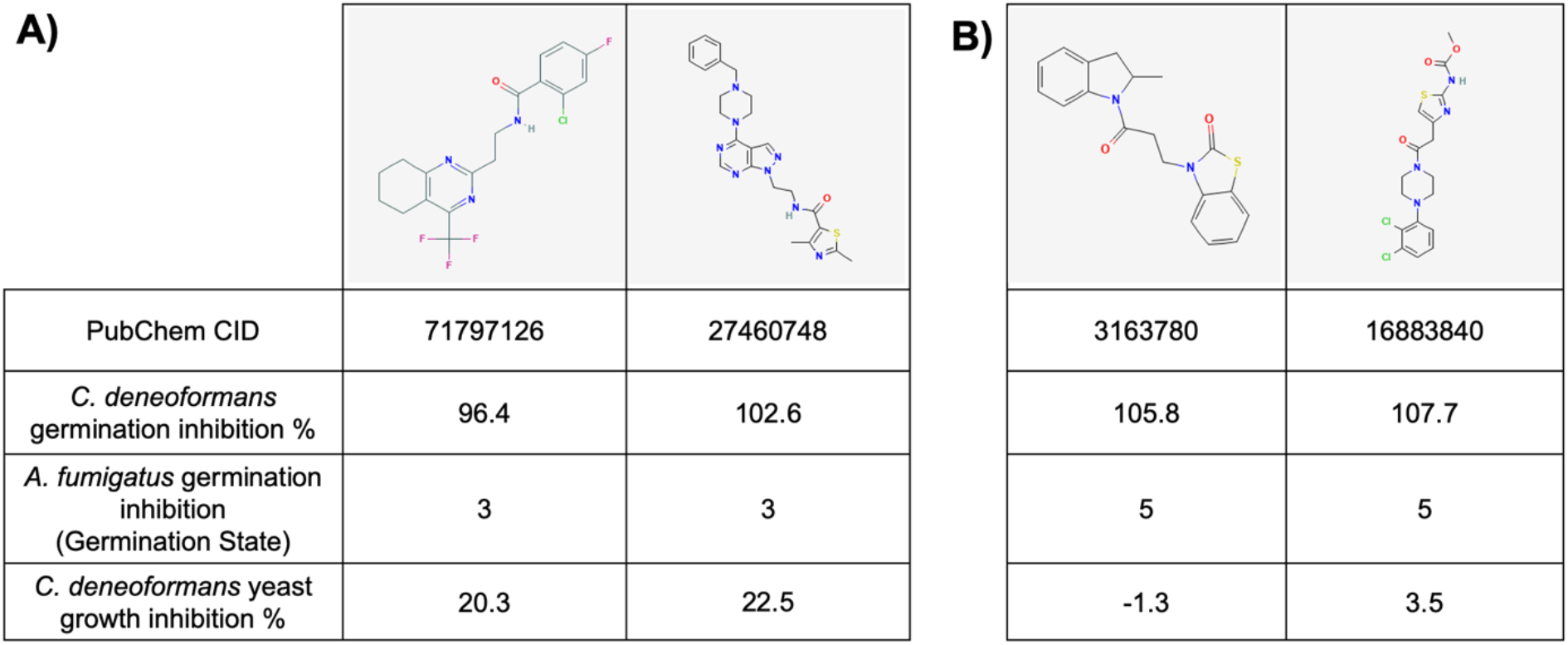
Identified exemplar compounds with research tool potential for A) identifying conserved molecular pathways broadly involved in fungal spore germination or B) identifying species-specific germination pathways. Exemplars were selected based on their abilities to selectively inhibit germination in *C. deneoformans* and *A. fumigatus* spore germination but not *C. deneoformans* yeast growth. Species-specific germination exemplars were identified by their abilities to inhibit spore germination only in *C. deneoformans. C. deneoformans* spore germination data are displayed as the percentage of inhibition as compared to a negative control in a nanoluciferase-based assay and *C. deneoformans* yeast growth data are displayed as percent growth inhibition compared to an untreated control (Ortiz *et al*., 2021). *A. fumigatus* conidia germination data are displayed as the Germination State identified in this work. Experiments were conducted with compounds in duplicate wells.

Within the identified group of germination-specific inhibitors, not all molecules inhibited germination in both *C. deneoformans* and *A. fumigatus*. There were 21 molecules that inhibited only *C. deneoformans* germination but not *A. fumigatus* (over 90% *C. deneoformans* germination inhibited, 0% *A. fumigatus* germination inhibited, and under 25% *C. deneoformans* yeast growth inhibited). For example, 3163780 and 16883840 (PubChem CIDs) showed nearly no inhibitory effects on *C. deneoformans* yeast growth and *A. fumigatus* germination but inhibited 100% of *C. deneoformans* germination **(Fig. 4B)**. Although the PubChem bioactivity database reveals no activity hits for compound 16883840, compound 3163780 has been marked as active in several screens of mammalian cells, including a screen for novel modulators of chloride ion-dependent transport processes and a screen for potassium channel inhibition (Delpire, PubChem Bioassay, Li *et al*., PubChem Bioassay). These results may suggest the involvement of compound 3163780 in transmembrane processes regulating cellular membrane potential, but further study is needed to confirm this speculated mechanism and its relevance to *C. deneoformans* germination.

To identify potential cellular targets of these compounds of interest (13 cross-species and 21 species-specific inhibitors), Swiss Target Prediction software was employed with default search parameters. There were no significant binding matches to potential targets for any of these molecules of interest, as the molecules either had a low binding probability to all proteins in the mammalian protein catalogs or only had a significant binding probability to mammalian proteins with minimal sequence similarity to any *C. deneoformans* or *A. fumigatus* protein. Therefore, the conserved or non-conserved targets of these molecules remain to be determined. Overall, these data suggest that some inhibitors are species-specific, whereas others are effective across species, which indicates potential for shared molecular targets that can be leveraged to learn more about fungal germination biology.

### A subset of shared germination inhibitors was also active against yeast growth in *Candida* species

Many identified *C. deneoformans* and *A. fumigatus* germination inhibitors also inhibit *C. deneoformans* yeast growth, which led us to ask whether these molecules could be active against two other priority fungal pathogens that grow primarily as yeast: *C. albicans* and *C. auris*.

To determine the efficacy of the 191 germination inhibitors against *C. albicans* (SC5314) and multi-drug-resistant *C. auris* (B11219) yeast growth, we carried out growth assays in the presence of each inhibitor. Briefly, *Candida* yeast were inoculated into liquid cultures containing 10 µM of each compound and assessed for growth using optical density (OD_600_). Five compounds inhibited growth in *C. albicans* with final optical densities at less than 80% of the no-compound (DMSO-only) controls **(Fig. 5A)**, and 20 compounds inhibited *C. auris* **(Fig. 5B)**. Four of these compounds inhibited yeast growth in both species **(Fig. 5C)**. Overall, we discovered that although *Candida* species are substantially phylogenetically diverged from *C. deneoformans*, some molecules were capable of cross-species yeast growth inhibition.

**Figure 5.**
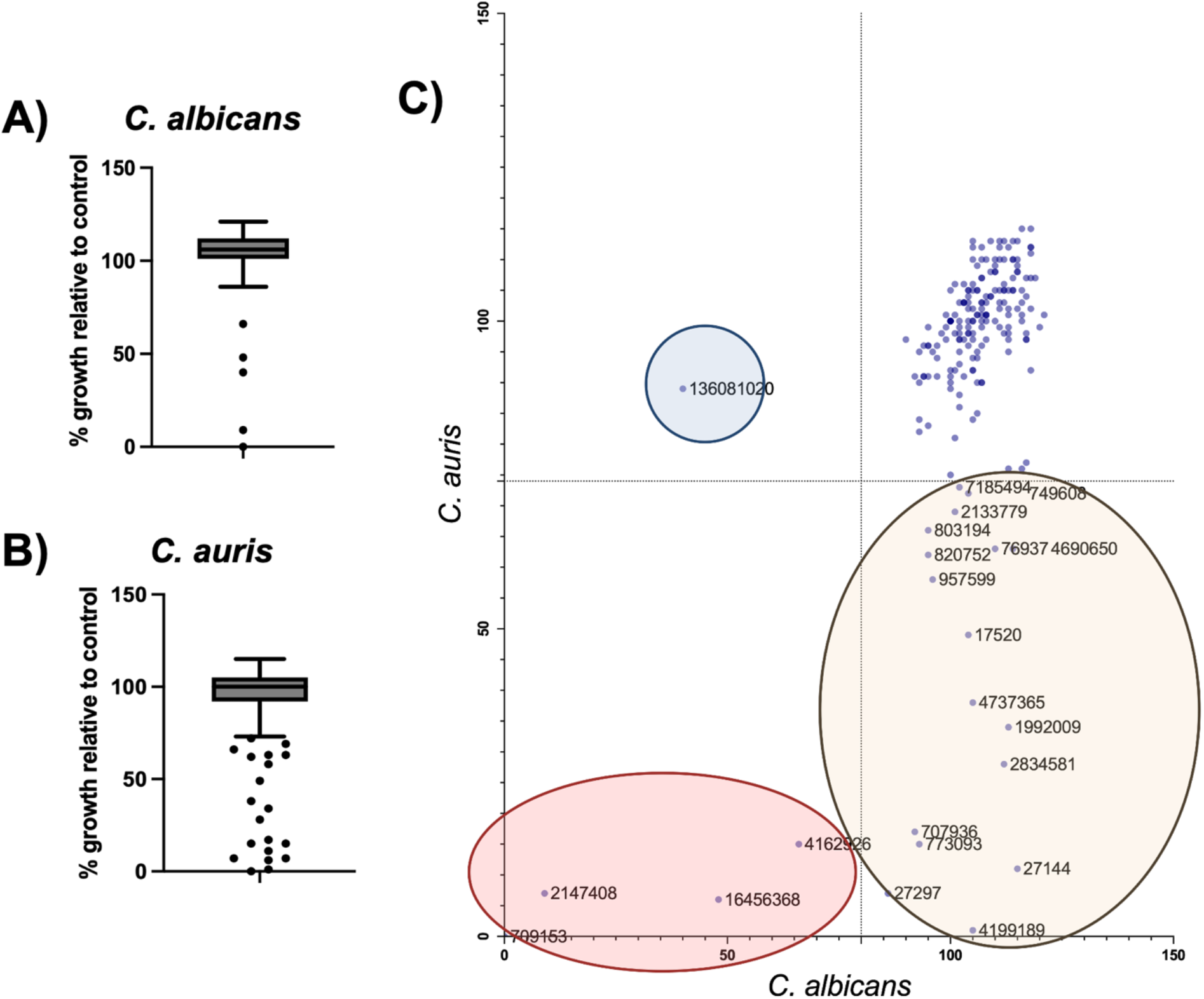
Inhibitor compounds were assessed for effects on *C. albicans* SC5314 and *C. auris* B11219 yeast growth at 10 µM. Box plots of percent growth relative to solvent (DMSO) control for each inhibitor compound at 24 hours for both **A)** *C. albicans* SC5314 and **B)** *C. auris* B11219. Relevant outliers are marked with dots. **C)** Percent growth relative to solvent control data is plotted to highlight the compounds uniquely inhibitory to *C. albicans* (blue oval), uniquely inhibitory to *C. auris*, (tan oval), and inhibitory to the growth of both species tested (pink oval). PubChem identifiers are shown for inhibitory compounds. Experiments were conducted with compounds in duplicate wells.

### A subset of the multi-species fungal inhibitors exhibits low cytotoxicity towards human cell types

Previous work revealed that 108 of the 191 originally identified *C. deneoformans* germination inhibitors are minimally cytotoxic to a cell line of normal human dermal fibroblasts (“minimally cytotoxic” defined as ≥ 80% cell viability) (Ortiz *et al*., 2021). To more broadly evaluate the toxicity of the fungal inhibitors, we tested two additional commonly used cell types: 1) human hepatocytes (HepG2 liver cancer cells) and 2) human erythrocytes (red blood cells). We used a commercial Nano Luciferase-based viability assay to assess cytotoxicity of the inhibitors at 10 µM toward hepatocytes and observed a wide range of viability, from inviable to completely viable. However, most compounds were minimally cytotoxic to hepatocytes at 10 µM with a median viability of 81.1% **(Fig. 6A)**.

**Figure 6.**
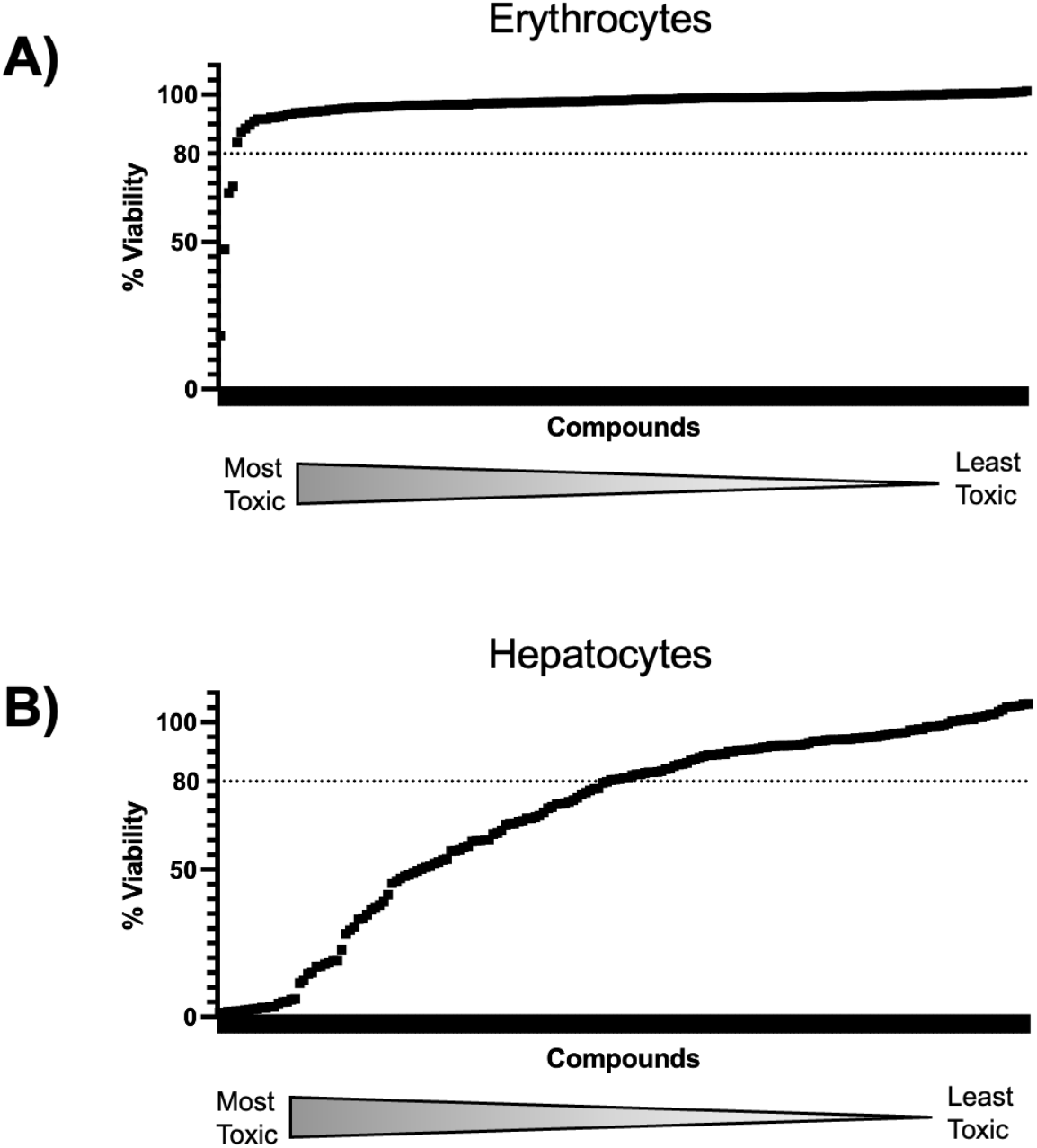
Inhibitor compounds were assessed for cytotoxic effects on human erythrocytes and hepatocytes. Dot plots show the percent viability of **A)** erythrocytes and **B)** hepatocytes (HepG2 cells) when treated with inhibitor compounds. Compounds that resulted in >80% viability when tested on either cell type were considered low toxicity. Triplicate wells of each compound were tested against both cell types. Viability data paired with compound ID is available in **Dataset S1**.

We determined erythrocyte viability in the presence of each molecule using a hemolysis assay, with higher OD_570_ measurements indicating higher hemolysis. Most inhibitor molecules were not toxic to erythrocytes at 10 µM, resulting in ~100% viability, with only four inhibitors as more hemolytic outliers with less than 80% cell viability **(Fig. 6B)**. Overall, a majority of the 191 fungal inhibitor molecules were minimally toxic to human cell lines (“minimal toxicity” defined as ≥ 80% cell viability), including hepatocytes and erythrocytes (with 100 and 187 inhibitors being minimally toxic, respectively).

### Nine broad spectrum fungal inhibitors with favorable properties for therapeutic development were identified

Data from screening experiments **(Fig. 2-6)** were compiled, normalized, and scored to create a list of compounds with the highest therapeutic potential **(Dataset S2)**. Among the top nine hits as determined by therapeutic potential score, three molecules (PubChem IDs 71781394, 72718563, and 71798856) shared similar substructures, all belonging to the previously defined “Substructure Group B” (B2, B8, and B5, respectively) **(Fig. 7A)** (Ortiz *et al*., 2021). Group B inhibitors share a substructure with a known class of agricultural fungicides thought to target succinate dehydrogenase in a fungal-specific manner (Sierotzki *et al*., 2013). These three molecules exhibited minimal cytotoxicity to the human cell lines tested and were strongly inhibitory against *C. deneoformans* germination and yeast growth as well as *A. fumigatus* germination. Although they did not inhibit growth of either *Candida* species, another molecule within the top hits, previously defined as a member of “Substructure Group D” (D14, PubChem ID: 2992099, shown in **Figure 7B**) was identified as a modest *C. auris* inhibitor and also exhibited moderate inhibitory effects on other fungal species tested. Overall, these molecules provide promising starting points for future development and optimization of antifungal compounds.

**Figure 7.**
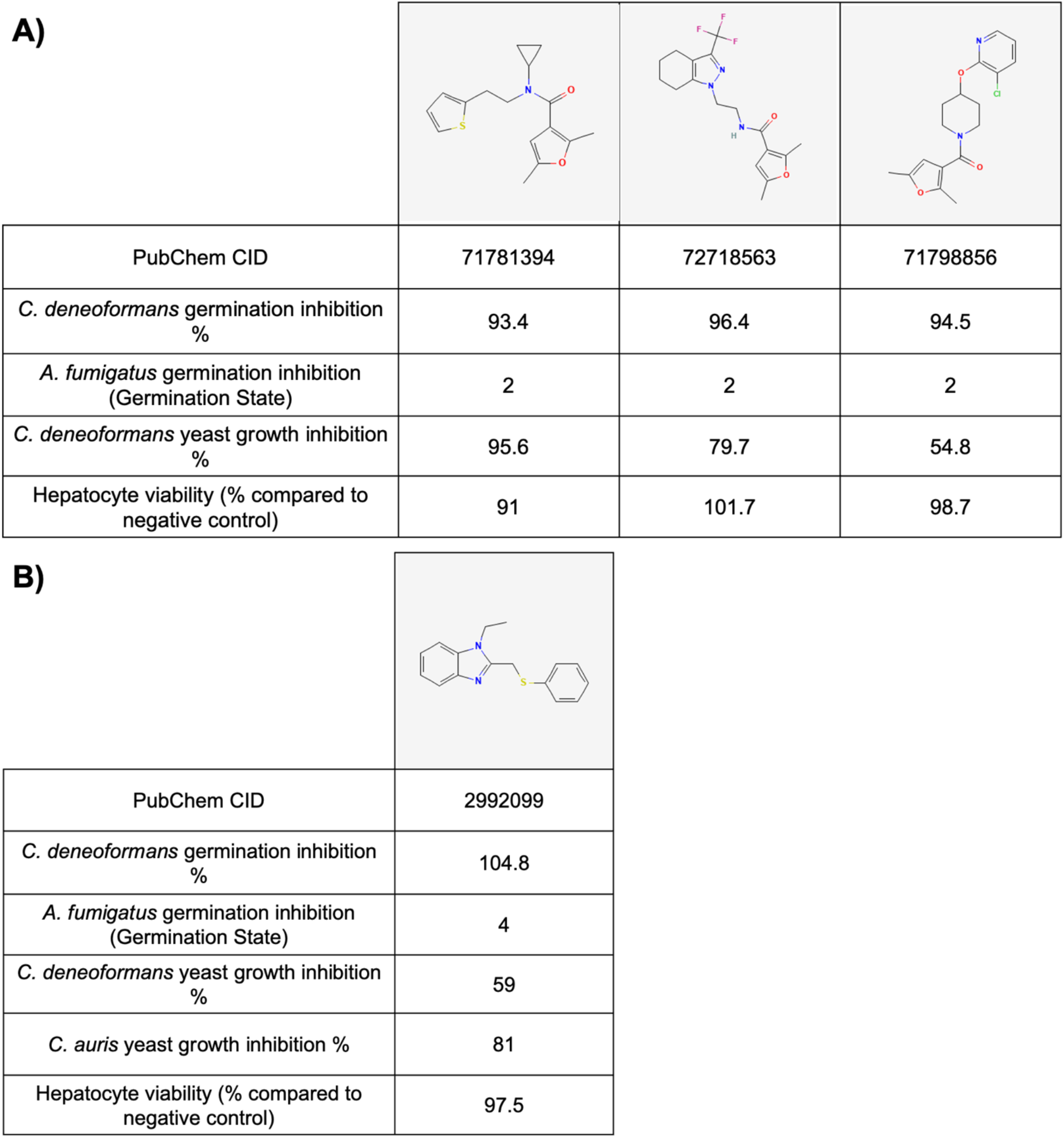
Identified exemplar compounds with therapeutic potential. **A)** Three members of Substructure Group B demonstrated favorable therapeutic properties (antifungal activity and nontoxicity to hepatocytes). **B)** A member of Substructure Group D also demonstrated favorable therapeutic properties including modest activity against *C. auris. C. deneoformans* spore germination data are displayed as the percentage of inhibition as compared to a negative control in a nanoluciferase-based assay and *C. deneoformans* yeast growth data are displayed as percent growth inhibition compared to an untreated control (Ortiz *et al*., 2021). *A. fumigatus* conidia germination data are displayed as the Germination State identified in this work. *C. auris* growth data displayed as percent of growth at 24 hours in the presence of the compound compared to a solvent (DMSO) control. Hepatocyte toxicity is given as the percent viability of HepG2 cells after incubation with the selected compound for 72 hours compared to solvent (DMSO) control. Experiments were conducted in either duplicate or triplicate as indicated previously.

None of these four exemplar compounds has been identified as hits in bioassays in the PubChem Database. However, using Swiss Target Prediction software on compounds of interest resulted in the prediction of potential targets of 2992099 and 71798856. The molecule 2992099 was predicted to have a 37.2% probability of binding a protein with 43% sequence identity to a histone deacetylase found in both *C. deneoformans* and *A. fumigatus*. The molecule 71798856 was predicted to have 24.3% probability of binding to an acetyl-CoA carboxylase (ACC) with sequence identities ranging from 29-37.2% to several biotin-dependent carboxylases in *C. deneoformans* and up to 46.4% sequence identity to ACC in *A. fumigatus*. This predicted target is metabolically connected with, but not structurally similar to, succinate dehydrogenase, the reported target of agricultural fungicides sharing a similar substructure (Sierotzki *et al*., 2013). More directly, ACC has been characterized as an essential enzyme in *Aspergillus nidulans*, a species related to *A. fumigatus*, and it has also been investigated as a potential drug target in *C. albicans* (Morrice *et al*., 1998) (Chen *et al*., 2023). Swiss Target Prediction was unable to identify likely targets for other molecules, but these existing predictions may guide future experimental efforts to confirm molecular targets of these identified fungal inhibitors.

## Discussion

In this study, we screened 191 previously identified *C. deneoformans* germination inhibitors for their abilities to inhibit germination and/or growth of other critical priority fungal pathogens. We discovered that 101 of these molecules inhibited *A. fumigatus* conidia germination, displaying endpoint phenotypes ranging from complete inhibition of germination to inhibition at different stages of conidial swelling or germ tube formation. We also discovered that five compounds inhibited *C. albicans* growth, 20 compounds inhibited *C. auris* growth, and four inhibited both *Candida* species. Of the 191 inhibitors, we found that 100 were minimally toxic (≥ 80% cell viability) to human erythrocytes and hepatocytes at concentrations inhibitory to fungi. Taking all these data together, we identified compounds with inhibitor/toxicity profiles that could be useful in the development of broad-spectrum antifungals or as research tools to identify germination-specific pathways and other potential molecular targets for inhibition.

Existing classes of antifungal drugs target cellular structures contributing to cell wall and membrane integrity, as well as nucleic acid synthesis (Mazu *et al*., 2016). Many of these drugs are known for their negative side effects in patients, which are caused by off-target activity on mammalian cell components. Spore germination, however, appears to be a fungal-specific process, so its associated targets warrant further investigation for the future development of lower-toxicity antifungal agents. With this motivation, we previously screened large libraries of small molecules (~75,000 compounds, Life Chemicals libraries 1-3) for inhibition of *C. deneoformans* germination (Ortiz *et al*., 2021). The molecular libraries used were comprised of structurally diverse small molecules with drug-like properties, including a design preference for low toxicity to human cells.

Given the phylogenetic distance between *C. deneoformans* and *A. fumigatus* (diverged at least 450 million years ago), we did not anticipate that the majority of our inhibitors of *C. deneoformans* germination would inhibit germination in *A. fumigatus* (Berbee *et al*., 2010). However, the data presented here highlight potential conservation of germination-related inhibitor targets between two species that undergo germination processes that appear quite morphologically distinct. This finding is promising because other spore-forming human fungal pathogens such as *Coccidioides immitis, Blastomyces dermatitidis*, and *Histoplasma capsulatum* are much more closely related to *A. fumigatus* and may also be susceptible to *C. deneoformans* germination inhibitors. An advantage of the *Cryptococcus* and *Aspergillus fumigatus* systems is that basidiospores and conidia from these fungi are Risk Group 2 (RG2) and can be handled with BSL2 precautions. If compounds that inhibit germination can be identified in these RG2 systems, then the more arduous task of testing against Risk Group 3 (RG3) spores (e.g. *C. immitis, B. dermatitidis, H. capsulatum*) can be limited to compounds with the highest likelihoods of efficacy.

Fifty-four of the compounds that inhibited *A. fumigatus* belonged to the previously identified structural groups, suggesting that germination mechanisms could be shared by phylogenetically distinct fungal pathogens. For example, there is evidence that inhibitors belonging to substructure Group B target mechanisms in oxidative phosphorylation in a fungal-specific manner to inhibit germination in *C. deneoformans*, and our study has revealed inhibition of *A. fumigatus* germination by Group B compounds as well (Ortiz *et al*., 2021). To define and characterize these conserved, “core” germination mechanisms, future work is needed to confirm the targets of broad-spectrum inhibitor compound groups.

These germination-specific inhibitors are of particular importance because they can be used to understand the molecular processes that distinguish germination from yeast growth. Because germination appears to be both spore-specific and required for pathogenesis, identifying germination-specific molecular targets promises to enable the development of broad spectrum, lower toxicity therapeutics. For example, the identified germination-specific inhibitors (exemplars highlighted in **Fig. 5**) could be useful tools for future studies to characterize the pathways required for germination and not vegetative yeast growth. In addition, the degree to which the molecular processes involved in spore germination are conserved among diverse species could provide an additional avenue for target discovery. For example, one group of identified inhibitors was active against both *C. deneoformans* and *A. fumigatus* germination, but others were active only against *C. deneoformans*. This finding highlights the presence of both conserved and species-specific germination pathways and provides opportunities to fine-tune small molecule inhibitors for specific fungi and create both broad-spectrum and pathogen-specific antifungal agents.

One of the most exciting uses of germination inhibitors clinically could be in the prevention of disease in vulnerable patient populations. Because individuals most susceptible to invasive fungal diseases are often severely immunocompromised, a highly effective, low toxicity antigerminant could be used prophylactically in vulnerable populations to prevent fatal disease. Furthermore, because fungal spores are a terminal cell type that cannot undergo vegetative growth without germinating first (fungal spores are not formed in the host), this type of antifungal drug might offer the advantage of circumventing the development of resistance by preventing the establishment of a resident replicating population in the host (Köhler *et al*., 2015, Fisher *et al*., 2012).

The work on *Candida* species in this study, specifically a pan-resistant *C. auris* isolate, addresses another way in which these molecules originally identified as germination inhibitors may aid in the ongoing antimicrobial resistance crisis. Although 101 inhibitors exhibited shared activity against both *C. deneoformans* and *A. fumigatus*, relatively fewer also had inhibitory activity towards *C. albicans* and *C. auris*. This may be because *Candida* species do not produce spores for infection and dispersal as *Cryptococcus* and *Aspergillus* species do, and these inhibitors were originally identified because of their germination inhibition abilities (Ortiz *et al*., 2021). However, we identified 20 compounds that inhibited yeast growth in a pan-antifungal drug-resistant *C. auris* strain (B11219). This finding reveals potential for new targetable pathway(s) to be identified in *C. auris* that may be optimized for the development of future therapeutics to combat existing and rising resistance in this species.

Importantly, we discovered that most molecules investigated in this work were minimally toxic to two human cell types (erythrocytes and HepG2 hepatocytes) in addition to the human dermal fibroblasts tested previously (Ortiz *et al*., 2021). Ultimately, the most promising molecules from these studies should be moved into preclinical development for investigation of long-term effects in cell culture and animal model assays. We hypothesize that germination-targeted therapeutics could help limit host toxicity due to the fungal-specific nature of spore and germination biology. Additionally, inhibitors with moderate therapeutic value scores have the potential to be chemically optimized to become even more effective antifungal agents with lower toxicity to mammalian cells.

## Methods

### *Aspergillus fumigatus* conidia germination assays

Glass-bottomed 384 well plates containing each *C. deneoformans* germination inhibitor in duplicate were acquired from the Small Molecule Screening Facility at the University of Wisconsin – Madison. The amount of inhibitor was added to each well such that the final concentration of inhibitor in the well during the experiment would be 10 µM. DMSO was used as a negative control. Plates were stored at −20°C until they were used.

*A. fumigatus* CEA10/A1163 was grown from frozen stock on solid glucose minimal media (GMM) for 3 days at 37°C. Conidia were collected into sterile water with 0.01% Tween 80 and ~16,000 conidia were added to each well in RPMI-MOPS pH 7. Plates were incubated on a Nikon Ti2 microscope set to 37°C and imaged every 2 hours for 8 hours. At endpoint, cells were visually inspected and assigned to Germination Groups 1-5, with 1 representing complete germination inhibition and 5 representing no difference from the DMSO control. *A. fumigatus* cell size and shape were quantified in ImageJ, then visualized as a two-dimensional histogram using a previously developed pipeline (Ortiz *et al*., 2024). For compounds that did not exhibit a germination inhibition phenotype at 10 µM, germination assays were performed again with each compound at final concentrations of 10, 20, 40, 80, and 100 µM in duplicate.

### *Candida* species growth assays

*Candida albicans* SC5314 and *Candida auris* B11219 were grown in overnight liquid yeast peptone dextrose (YPD) cultures. Cells were enumerated by hemocytometer and resuspended to 5 × 10^5^ cells/mL in YPD. Cells were added to 96-well plates pre-loaded with inhibitor compounds in duplicate wells at a concentration of 5 × 10^4^ yeast cells per well at a total end volume of 100 µL per well (OD_600_~ 0.005) and inhibitor concentration of 10 µM. Plates were incubated at 30°C, stationary. At 0, 6, and 24 hours, plates were mixed by pipetting, and OD_600_ was recorded. Data were analyzed in GraphPad Prism using ROUT (robust regression and outlier removal) outlier analysis using both Q = 5% and Q = 10%.

### Hepatocyte cytotoxicity assay

HepG2 cells were added to Corning 3765 well plates using a Multidrop Reagent Dispenser (Thermo Fisher) in 50 µL of RPMI medium and allowed to attach overnight at 37°C. Inhibitor molecules were then added in triplicate wells using the Echo Acoustic Dispenser (LabCyte). Doxorubicin and DMSO at 10 µM were used as positive and negative controls respectively. Following 72 hours of incubation, cell viability was assayed via CellTiter-Glo (Promega). 25 µL of CellTiter-Glo reagent was added per well. Following ten minutes of incubation, luminescent signal was detected using a Pherostar plate reader (BMG). Toxicity was calculated by determining the percent reduction in signal as compared to the DMSO negative control.

### Erythrocyte hemolysis assay

Inhibitor molecules were added to 384 well plates (Thermo Fisher 264574) in triplicate using the ECHO 550 (Labcyte) for a final concentration of 10 µM. 0.1% Triton-X was used as a positive hemolysis control. Human erythrocytes (BioIVT) were washed with PBS and diluted to a concentration of 6 × 10^7^ cells/mL. 50 µL were incubated with inhibitor molecules using the Biomek FX (Beckman) for one hour shaking at room temperature and subsequently spun down at 4000 RPM for 10 minutes to pellet cells. 30 µL supernatant was removed and added into a 384 well clear plate (Phoenix Scientific MPG-781186) using the Biomek FX. Absorbance was read at OD_570_to quantify hemolysis, with higher ODs indicative of hemolytic activity.

### Classification of molecules as inhibitors

Collected data was compiled into a spreadsheet, and weighted, optimized scores were generated to rank molecules for germination-specific inhibition (both cross-species and *C. deneoformans*-specific) and therapeutic potential **(Dataset S2)**. Assay results were weighted linearly based on molecule concentration, and when applicable, safety data (human cell viability) was weighted 60% and efficacy data (antifungal activity) was weighted 40%. Thresholds were set to bin molecules into inhibitor groups and are defined in the results section.

## Acknowledgements

We thank the Small Molecule Screening Facility at the University of Wisconsin-Madison Carbone Cancer Center supported by P30 CA014520, including Dr. Spencer Ericksen and Song Guo, for their assistance and expertise preparing screening assay plates and conducting mammalian cell cytotoxicity assays. We thank Harrison Estes for providing *A. fumigatus* conidia for experiments. We thank Dr. Soleil Young for cladogram construction. We also thank Nicolas Pereira, Sehrish Afsheen, Samantha McKechnie, and Kamea Teller for their comments on the manuscript.

J.A.S. was supported in part by an NIH T32 award to the UW Chemistry–Biology Interface Training Program (T32GM008505). This work was supported by NIH R01 AI179964 to C.M.H. The content is solely the responsibility of the authors and does not necessarily represent the official views of the National Institutes of Health.

